# Range of *Ixodes laguri*, a nidicolous tick that parasitizes critically endangered rodents, with details on its western distribution limit in Austria

**DOI:** 10.1101/2024.02.20.581178

**Authors:** Franz Rubel

## Abstract

The nidicolous tick *Ixodes laguri* is a nest-dwelling parasite of small mammals that mainly infest rodents of the families Cricetidae, Gliridae, Muridae and Sciuridae. There is no proven vectorial role for *I. laguri*, although it is suggested that it is a vector of *Francisella tularensis*. In this study, a first map depicting the entire geographical distribution of *I. laguri* based on geo-referenced locations is presented. For this purpose, a digital data set of 141 georeferenced locations from 16 countries was compiled. Particular attention is paid to the description of the westernmost record of *I. laguri* in the city of Vienna, Austria. There, *I. laguri* is specifically associated with its main hosts, the critically endangered European hamster (*Cricetus cricetus*) and the European ground squirrel (*Spermophilus citellus*). These two host species have also been mapped in the present paper to estimate the potential distribution of *I. laguri* in the Vienna metropolitan region. The range of *I. laguri* extends between 16–108° E and 38–54° N, i.e. from Vienna in the east of Austria to Ulaanbaatar, the capital of Mongolia. In contrast to tick species that are expanding their range and are also becoming more abundant as a result of global warming, *I. laguri* has become increasingly rare throughout its range. However, *I. laguri* is not threatened by climate change, but by anthropogenic influences on its hosts and their habitats, which are typically open grasslands and steppes. Rural habitats are threatened by the intensification of agriculture and semi-urban habitats are increasingly being destroyed by urban development.

## 1. Introduction

The geographical distribution of the nidicolous tick *Ixodes laguri* Olenev, 1929 is only partially known. So far, two maps have been published showing georeferenced locations of *I. laguri*. The first map by Feider (1965) shows 13 locations between Hungary in the west and Kazakhstan, the region north of the Aral Sea, in the east. The second, more up-to-date map by Mihalca and D’Amico (2017) shows three times as many locations, but goes only as far as to Ukraine and Turkey in the east. A further map, published online by Kolonin (2009), did not contain any georeferenced tick locations but estimated the range of *I. laguri* as described by Siuda and Sebesta (1997). Unfortunately, this map is no longer online and was not available for this work either. Reference is therefore made to the earlier map by Kolonin (1981). In summary, there is no map that covers the entire known distribution area of *I. laguri* based on georeferenced findings. However, there is a current list on the geographic distribution of *I. laguri* by countries and territories (Guglielmone et al., 2023).

*Ixodes laguri* is a nest-dwelling parasite of small mammals that mainly infests rodents of the families Cricetidae, Gliridae, Muridae and Sciuridae. The most comprehensive list of all host animals on which *I. laguri* has been found was compiled by Anastos (1957). Cases of people being attacked by immature stages have been reported by Filippova (1977), and female adult *I. laguri* found on humans have been reported from Turkey (Bursali et al., 2011; Keskin et al., 2015). It is assumed that there is no difference in the host preferences of immatures and adults, with the exception that adult males are not parasitic (Siuda and Sebesta, 1997). As a very host-specific tick, *I. laguri* is only found in the habitats of its hosts as mentioned above. In Central Europe, the main hosts are European hamster (*Cricetus cricetus*) and European ground squirrel (*Spermophilus citellus*), which inhabit cultivated lands such as fallows, grasslands, vineyards, but also golf courses and airfields (Rubel, 2024). Further east, *I. laguri* also inhabits steppe, less often semisteppe and semidesert habitats. In Trans-Caucasia, i.e. Armenia, Georgia, and Azerbaijan, the tick inhabits mountain steppes up to altitudes of 1500 m, where Kirschenblatt (1936) found it on gerbils (*Meriones tristrami*), steppe lemmings (*Lagurus lagurus*), hamsters (*C. cricetus*, *Mesocricetus brandti*) and ground squirrels (*Spermophilus pygmaeus*).

The three-host tick *I. laguri* is nidicolous (Gray et al., 2014), although it has occasionally been found questing in very close proximity to the burrows of its hosts (Mihalca and D’Amico, 2017). Usually only a few tick specimens were found in individual nests. A maximum of 28 larvae and 100 nymphs was collected from one nest of the European ground squirrel by C^̌^erný (1990). In the 1970s, Honźaková et al. (1980) investigated the development of *I. laguri* in the nests of the European ground squirrel. For this purpose, ticks were taken from the nests of wild ground squirrels in Slovakia and kept in a field experiment as well as under controlled laboratory conditions. The measurements of the microclimate in ground squirrel nests previously carried out in the field showed an average temperature of 15-17° C. In a burrow depth of 2 m, these temperatures occur in June–October. Additionally, a high relative humidity of 90% was measured. The application of these rather constant conditions in the laboratory experiments, together with the field experiments, led to the determination of the natural life cycle of *I. laguri* of 2–3 years. Life cycles significantly less than one year, as determined by Russian authors (Anastos, 1957), are not possible in Central Europe and the Balkans. Honźaková et al. (1980) describes the following life cycle in Slovakia: Females moult in mid-summer, feed in April of the following year and oviposit in May. Larvae hatch and feed in August. Nymphs moult in November, hibernate in a hungry state and feed in April. Adults moult in July. This is in accordance with observations in Romania, where the highest female activity has been reported in spring and larvae were found in summer. Peaks in nymphal *I. laguri* activity have been reported in spring and autumn (Feider, 1965).

Adequate illustrations and morphological characteristics of *I. laguri* can be found in Feider (1965) and Mihalca and D’Amico (2017), although the four subspecies introduced in the historical literature (Anastos, 1957) are not discussed. Biometric measures of morphological features compared to *Ixodes ricinus*, *Ixodes persulcatus* and *Ixodes acuminatus* (syn. *I. redikorzevi*) have been investigated by Voltzit and Pavlinov (1996). Haller’s organ of *I. laguri* was also examined using scanning electron microscopy (Honzáková et al., 1975; Suppan, 2013). The 16S rDNA (Orkun, 2018), COX1 (Radulovíc et al., 2017; Numan et al., 2023), and ITS2 (Radulovíc et al., 2017) phylogenetics indicate that *I. laguri* is a member of the *I. ricinus* complex.

The only evidence of pathogens occurring in *I. laguri* dates back to work in the former Soviet Union in the 1950s. Accordingly, the causative agent of tularaemia, *Francisella tularensis*, was detected in *I. laguri* in Volgograd Oblast and infection with rickettsia was also reported (Zhmaeva and Korshunova, 1953; Philip and Burgdorfera, 1961; Pollitzer, 1963). Based on this and the fact that *I. laguri* has often been found in tularaemia endemic areas (Bozhenko and Shevchenko, 1956), the tick has repeatedly been referred to as a vector of *F. tularensis* (Kiefer et al., 2010; Musaev et al., 2019; Zabashta et al., 2022). However, without proven capability of transmission, the vector function of a given tick species for a given pathogen is not substantiated (Kahl et al., 2002). Since there are no transmission experiments known with *I. laguri* for any pathogen, no vector function has been proven.

## 2. Materials and Methods

To determine the range of *I. laguri* in Eurasia, the literature list of reviews on the geographical distribution of *I. laguri* (Siuda and Sebesta, 1997; Guglielmone et al., 2023) was used and supplemented by further literature covering the western distribution area. Since the publications on documented localities of *I. laguri* in its eastern distribution area are not listed in PubMed or Scopus, an extensive Google and Cyber Leninka search was carried out in Russian and Ukrainian language. In addition, the translated summary of the United States Department of Agriculture (USDA) was used, in which historical *I. laguri* locations from Azerbaijan, Bulgaria, Georgia and Russia were documented (Doss et al., 1978). According to Table 1 the following numbers of *I. laguri* locations were incorporated: 4 in Armenia, 1 in Austria, 4 in Azerbaijan, 5 in Bulgaria, 4 in Georgia, 3 in Hungary, 15 in Kazakhstan, 2 in Moldova, 2 in Mongolia, 7 in Romania, 36 in Russia, 11 in Serbia, 18 in Slovakia, 14 in Turkey, 1 in Turkmenistan, and 14 in Ukraine.

**Table 1:** Number, accuracy (high, medium, low and very low), country, and reference of georeferenced *Ixodes laguri* sampling sites compiled in this study.

| No. | Acc. | Country | References |
| --- | --- | --- | --- |
| 1 | v | Armenia | Anastos (1957) |
| 1 | l | Armenia | Filippova (2008) |
| 2 | l | Armenia | Dilbaryan and Hovhannisyan (2016) |
| 1 | h | Austria | Suppan (2013) |
| 1 | v | Azerbaijan | Anastos (1957) |
| 3 | v | Azerbaijan | Doss et al. (1978) |
| 2 | v | Bulgaria | Doss et al. (1978) |
| 1 | l | Bulgaria | Hubálek (2009) |
| 2 | l | Bulgaria | Arnaudov and Georgiev (2022) |
| 2 | v | Georgia | Doss et al. (1978) |
| 2 | l | Georgia | Shaposhnikova and Sakhno (2012) |
| 3 | l | Hungary | Janisch (1959) |
| 5 | v | Kazakhstan | Anastos (1957) |
| 1 | v | Kazakhstan | Feider (1965) |
| 4 | v | Kazakhstan | Doss et al. (1978) |
| 3 | l | Kazakhstan | Amirova et al. (1989) |
| 1 | l | Kazakhstan | Katuova et al. (2021) |
| 1 | v | Kazakhstan | Kdyrsikh et al. (2021) |
| 1 | v | Moldova | Feider (1965) |
| 1 | m | Moldova | Vladimirovna (2014) |
| 2 | l | Mongolia | Kiefer et al. (2010) |
| 2 | v | Romania | Feider (1965) |
| 5 | l | Romania | Mihalca et al. (2012) |
| 7 | v | Russia | Anastos (1957) |
| 3 | v | Russia | Doss et al. (1978) |
| 13 | l | Russia | Filippova and Stekolnikov (2007) |
| 1 | l | Russia | Filippova (2008) |
| 1 | v | Russia | Kormilenko and Moskvitina (2009) |
| 1 | v | Russia | Lesheva et al. (2013) |
| 1 | v | Russia | Obert et al. (2015) |
| 3 | l | Russia | Grigor'evich (2016) |
| 2 | l | Russia | Kirillova and Kirillov (2018) |
| 1 | v | Russia | Musaev et al. (2019) |
| 3 | l | Russia | Porshakov et al. (2020) |
| 11 | l | Serbia | Radulović et al. (2017) |

Table 1: (continued)
| No. | Acc. | Country | References |
| --- | --- | --- | --- |
| 18 | l | Slovakia | Černý (1990) |
| 3 | l | Turkey | Arthur (1957) |
| 3 | l | Turkey | Keskin et al. (2007, 2015, 2019) |
| 6 | v | Turkey | Bursali et al. (2011, 2015) |
| 2 | l | Turkey | Güven et al. (2022) |
| 1 | l | Turkmenistan | Berdyayev and Annayev (1997) |
| 1 | v | Ukraine | Anastos (1957) |
| 1 | m | Ukraine | Stupnitskaya et al. (1964) |
| 1 | v | Ukraine | Feider (1965) |
| 2 | v | Ukraine | Sljar (1970) |
| 2 | v | Ukraine | Sklyar (2002) |
| 1 | v | Ukraine | Kuznetsov and Bondarev (2007) |
| 4 | l | Ukraine | Voronova et al. (2012) |
| 1 | l | Ukraine | Akimov and Nebogatkin (2016) |
| 1 | l | Ukraine | Evstafiev (2017) |
| **141** |  | **Total** |  |

Almost all tick findings were digitized based on text information on the locations or printed maps. These digitized locations, of course, are generally of lower accuracy than locations described by geographical coordinates determined by GPS in the field. To provide evidence of this, accuracy measures were given for all data referenced in Table 1 in accordance with a scheme established in previous studies (Rubel and Brugger, 2022). It is distinguished between high (h ≈ ±0.1 km), medium (m ≈ ±1 km), low (l ≈ ±10 km) and very low (v) accuracies. The latter has been applied here for the first time, and identifies all reports that relate only to political districts, mountain ranges or river sections. Such information is obligatory in Russian and Ukrainian literature and was used in the absence of more precise location information.

Despite including all available information, the number of georeferenced locations of *I. laguri* is one order of magnitude lower than that of comparable tick species such as *Ixodes trianguliceps* (Rubel and Kahl, 2023). This is also because *I. laguri* is associated to specific hosts, some of which are now threatened with extinction. The distribution of the hosts is therefore of particular importance, as it can indirectly be used to determine the distribution of *I. laguri*. The main host species were determined using an abundance count. For the metropolitan area of Vienna, the distribution map of main hosts, i.e. European hamsters and European ground squirrels, from Rubel (2024) was adapted to depict the exact locality of *I. laguri*. For this purpose and to visualize the global distribution of *I. laguri*, the georeferenced locations were plotted on terrain maps (OpenStreetMap contributors, 2017).

## 3. Results and Discussion

The main result of the present study is a map of the entire distribution areas of the tick *I. laguri* with the total number of 141 georeferenced locations for the period 1957–2022 (Fig. 1). The westernmost location was documented in the city of Vienna, Austria, at about 16° E (Fig. 2), and is confirmed by a location in Dvorniky, Slovakia, 100 km further to the east (C^̌^erný, 1990). The eastern distribution limit, however, is considered less certain. It may be at 82° E, where several authors have documented locations on both sides of the Russia-Kazakhstan border (Amirova et al., 1989; Obert et al., 2015; Grigor’evich, 2016). However, two locations were also reported near the Mongolian capital Ulaanbaatar, which is 1900 km further east. The first record of *I. laguri* in Mongolia dates back to 1974 (Dash, 1986), after which the tick was found on the Mongolian gerbil (*Meriones unguiculatus*) 10 km southeast of Ulaanbaatar. In 2006, the same German-Mongolian consortium documented the tick northwest of Ulaanbaatar, also on Mongolian gerbils (Kiefer et al., 2010). These findings are considered realistic because the type of habitat and the host species are similar to those in the western distribution area, and Josef Nosek, a recognized expert in the morphology of ticks from the Slovak Academy of Science (Nosek and Sixl, 1972), was involved in the first description of this tick species in Mongolia. However, to date, confirmation of the occurrence of *I. laguri* in Mongolia by other independent scientists is still pending. The eastern distribution limit was therefore provisionally set at 108° E. The north-south extension is largest in the Asian part of the range and extends from the steppe of Turkmenistan at about 38° N (Berdyyev and Annayev, 1997) to Western Siberia, Russia, at about 54° N (Grigor’evich, 2016). Note that only a single record of *I. laguri* has been found in Turkmenistan, so this southernmost record also awaits confirmation. It is also worth mentioning that in Central Europe *I. laguri* generally only occurs south of 49° N. The distribution area therefore extends between 16–108° E and 38–54° N, i.e. from Vienna in the east of Austria to Ulaanbaatar, the capital of Mongolia.

**Figure 1:**
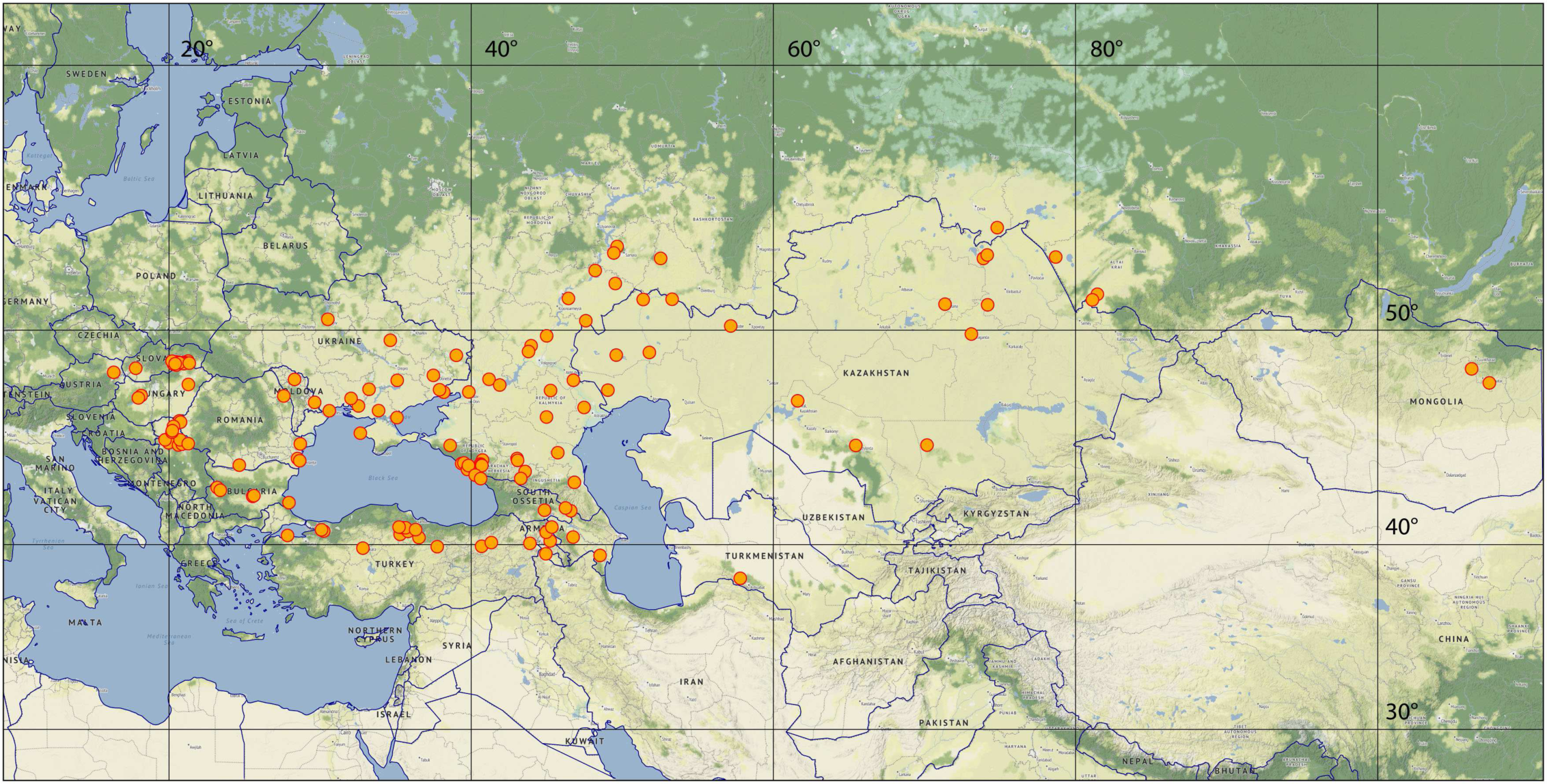
Findings of *Ixodes laguri* (orange points) in open grasslands, steppes and mountain steppes ranging between 16–108° E and 38–54° N.

**Figure 2:**
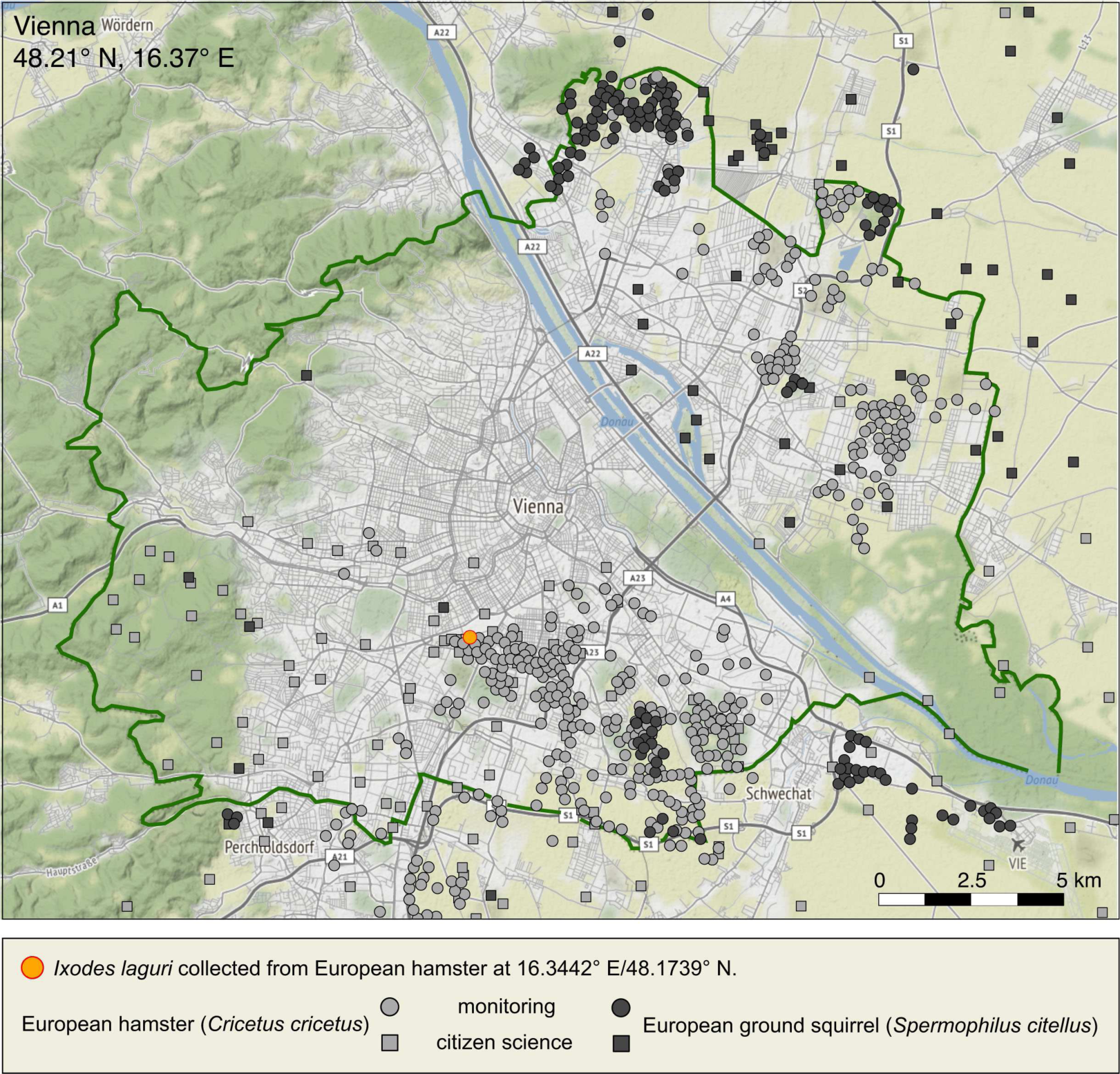
City map of Vienna showing the exact location of *Ixodes laguri* together with the distribution of its main hosts, the European hamster (*Cricetus cricetus*) and the European ground squirrel (*Spermophilus citellus*). Host data representative for the period 2000–2023 according to Rubel (2024).

A frequency distribution of *I. laguri* hosts is shown in Fig. 3. If one takes into account only those studies from which georeferenced locations could be generated (Table 1), then most of the studies report the tick *I. laguri* on hamsters (*C. cricetus*, *Nothocricetulus migratorius*, *Mesocricetus auratus*, *Mesocricetus newtoni*, *Mesocricetus raddei*, *M. brandti*) and ground squirrels (*S. citellus*, *S. pygmaeus*). Other hosts, summarized in the frequency distribution in Fig. 3, are voles (*Arvicola terrestris*, *Myodes rutilus*, *Myodes glareolus*, *Microtus subterraneus*), mice (*Mus musculus*, *Apodemus agrarius*, *Apodemus sylvaticus*), gerbils (*M. unguiculatus*, *M. tristrami*), hedgehogs (*Erinaceus concolor*), rats (*Rattus norvegicus*, *Hemiechinus auritus*), moles (*Talpa altaica*), blind mole-rats (*Spallax* sp.), shrews (*Crocidura* sp.), marmots (*Marmota sibirica*), and lemmings (*L. lagurus*). In addition, the tick was found in Ukraine on predators such as the steppe polecat (*Mustela eversmanii*) and the red fox (*Vulpes vulpes*) (Sklyar, 2002).

**Figure 3:**
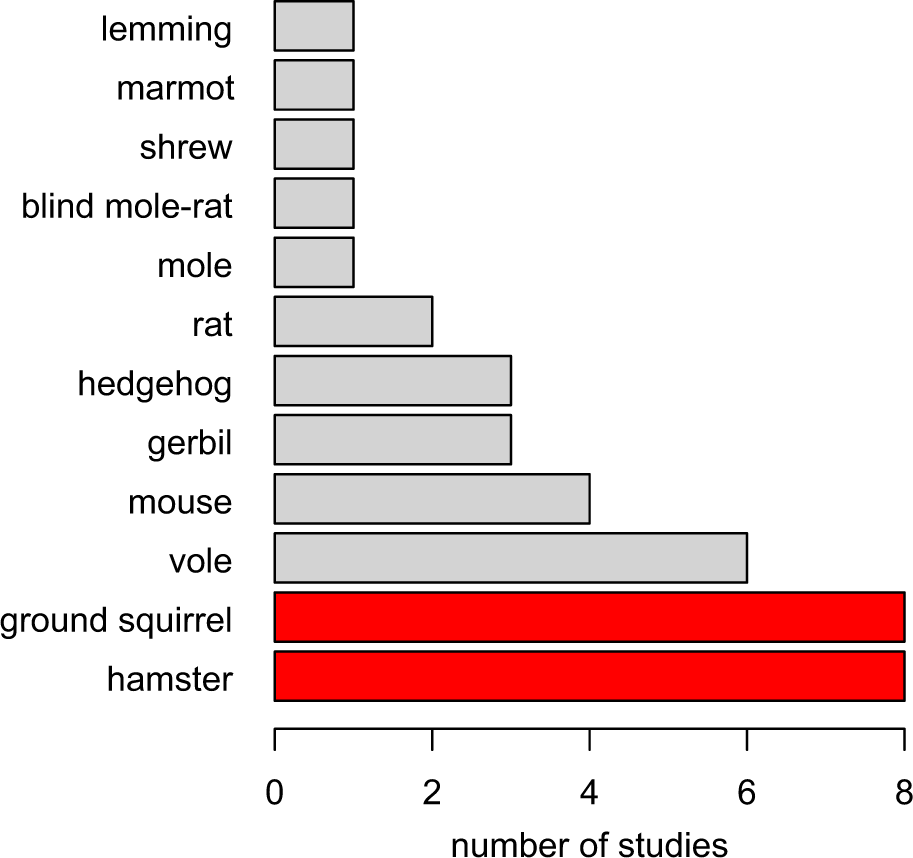
Number of studies reporting locations of small mammals invested with *Ixodes laguri*. Most studies (red bars) report finding the tick on hamsters (mainly *Cricetus cricetus*) and ground squirrels (mainly *Spermophilus citellus*).

Although the occurrence of *I. laguri* in Austria was already mentioned by Radda et al. (1986), there is currently no description of a location in the international literature, which is now being done here. Fig. 2 depicts a recent location of *I. laguri* in the city of Vienna, Austria. These specimens of *I. laguri* were collected from European hamsters, which were caught with live traps on the grounds of the clinic Favoriten at the geographical coordinate 16.3442° E/48.1739° N (Siutz and Millesi, 2012). In this study from 2005, reproductive timing in female hamsters, which affects offspring development and survival, was investigated. Therefore, the study by Siutz and Millesi (2012) does not contain any information about collected ticks, although these were mentioned in the master thesis of Suppan (2013). The latter examined the structure of dermal glands associated with spiracles using a scanning electron microscope, primarily of *I. ricinus* but also of *I. laguri*. Since there are no further data on the occurrence of *I. laguri* in Vienna, this location was overlaid on a map of the most important hosts, the European hamster (*C. cricetus*) and the European ground squirrel (*S. citellus*), which was adapted from Rubel (2024). From this map, at least the possible range of *I. laguri* in Vienna can be estimated, because ticks specifically associated with their hosts are co-distributed with them (Mihalca et al., 2012). For example, the distribution area of *I. laguri* from eastern Austria and Slovakia across the Balkans to the Black Sea (Fig. 1) corresponds almost exactly to the global distribution of *S. citellus* (Ramos-Lara et al., 2014).

In the metropolitan area of Vienna, which includes the outskirts of the city (Fig. 2), the population of hamsters can be estimated at 4,000 and that of ground squirrels at 16,000 individuals (Rubel, 2024). Thus, the total number of these main hosts of *I. laguri* in the metropolitan area of Vienna is about 20,000 individuals. The systematic examination of European ground squirrels for parasitic ticks in Serbia, 500 km away, indicates that *I. laguri* may not be a rare tick species in Vienna. Radulovíc et al. (2017) found more than 1,000 ticks on 151 infested ground squirrels in this study, with 79% identified as *I. laguri* and 21% as *Haemaphysalis concinna*. Ixodid ticks of the genera *Ixodes* and *Haemaphysalis* were also found on European hamsters in Turkey (Uslu et al., 2008), where *I. laguri* in rare cases also infests humans (Keskin et al., 2007, 2015). In the Caucasus, Filippova and Stekolnikov (2007) found larvae of *I. laguri* together with larvae of *I. ricinus* and *D. reticulatus*. *Ixodes laguri* (Suppan, 2013) as well as *I. ricinus* and *D. reticulatus* (Vogelgesang et al., 2020; Rubel and Brugger, 2022) also occur sympatrically in Vienna. That there is only one documented location of *I. laguri* in Vienna up to now is probably due to the fact that no studies concerning ticks on small mammals were carried out for a long time. Since 2019, *C. cricetus* and *S. citellus* are listed on *The IUCN Red List of Threatened Species* as critically endangered (Banaszek et al., 2020; Hegyeli, 2020). Investigations of *I. laguri* in the nests of *S. citellus*, like those from the field expeditions 1959–1976 (C^̌^erný, 1990), are therefore hardly possible anymore in the European Union.

Further, a sharp decline in the main host populations of *I. laguri* has been observed over the last few decades. For example, the global *C. cricetus* population has declined by 75% (Surov et al., 2016). Something similar has been reported for the Ciscaucasian or Georgian hamster (*M. raddei*), whose population has also declined massively. Responsible for this were the widespread ploughing of virgin and fallow lands, which began in the middle of the last century, and the reduction of natural pasture areas. As a consequence, this also led to a decrease of *I. laguri*, the suspected main vector of the causative agent of tularaemia in that region (Zabashta et al., 2022). There is strong evidence that *I. laguri* is one of those tick species that are becoming rarer along with their hosts.

In summary, the 141 georeferenced *I. laguri* locations come from the following 16 countries: Armenia, Austria, Azerbaijan, Bulgaria, Georgia, Hungary, Kazakhstan, Moldova, Mongolia, Romania, Russia, Serbia, Slovakia, Turkey, Turkmenistan, and Ukraine. For comparison, Guglielmone et al. (2023) listed 15 countries. Their list also includes the Czech Republic, for which no data is available since all of locations described in former Czechoslo-vakia (C^̌^erný, 1990) are in today’s Slovakia. Conversely, the locations in Serbia (Radulovíc et al., 2017) and the location in Austria described here are missing from the list of countries by Guglielmone et al. (2023). The latter was missing because there was no reference in the international literature to date. In the book chapter on the distribution of *I. laguri* by Mihalca and D’Amico (2017), the occurrence in Serbia (Radulovíc et al., 2017) could not be taken into account because it was published at the same time. In addition, the occurrence of *I. laguri* in Belarus and the Baltic countries Estonia, Latvia and Lithuania mentioned in the text might be an error because these countries are too far north and no locations are depicted even in the map of Mihalca and D’Amico (2017). This mismatch has already been considered by Guglielmone et al. (2023). The Republic of Dagestan should also not be listed as a separate country because it has been part of Russia since 1991. No data could be found for Uzbekistan either, which is why the work of Mihalca and D’Amico (2017) should be revised before it is used as a reference for the distribution of *I. laguri*. In contrast, the distribution of *I. laguri* described in the review by Siuda and Sebesta (1997) largely agrees with the collection of georeferenced locations presented here. Accordingly, the distribution area west of the Caspian Sea is divided into a wider northern and a narrower southern part. The northern part extends from the southern Lower Volga areas via Kazakhstan into Mongolia, while the southern part extends through the Caucasus and Trans-Caucasia to western Turkmenistan.

## 4. Conclusions

Based on georeferenced locations, the first map of *I. laguri* covering the entire distribution area was compiled. For this purpose, the westernmost location of *I. laguri* in the city of Vienna was mapped along with its main hosts, hamsters and ground squirrels. The findings at the eastern distribution limit in Ulaanbaatar, Mongolia, were also critically discussed. In addition, a list of 16 countries where the tick has been reported was compiled. However, it must be noted that knowledge about the range, biology and vector competence of *I. laguri* is limited, especially in comparison to other much more prominent tick species of the *I. ricinus* complex (Kahl and Gray, 2023). Since *I. laguri* can only be found on its often highly endangered hosts and in their nests, it can be assumed that this will remain the case in the future. In contrast to tick species that are expanding their range and are also becoming more abundant as a result of global warming, such as *I. ricinus* (Jaenson et al., 2012; Nuttall, 2021) and *D. reticulatus* (Mierzejewska et al., 2015; Brugger and Rubel, 2023), *I. laguri* is becoming increasingly rare. *Ixodes laguri*, a nidicolous tick of grasslands and steppes, is not threatened by climate change, but by anthropogenic influences on its hosts and their habitat. Rural habitats are threatened by the intensification of agriculture and semi-urban habitats are increasingly being destroyed by urban development (Rubel, 2024).

